# On the variability of Species Abundance Distributions with trophic guild and community structure

**DOI:** 10.1101/289348

**Authors:** David García-Callejas

## Abstract

Species Abundance Distributions (SADs) are one of the strongest generalizations in community ecology, but their variation across trophic levels remains largely unexplored. I study the variation in SAD metrics across trophic guilds in model and empirical communities. First, I use a theoretical model that allows tracking the variations in abundances across trophic levels, accounting for variations in species vulnerability. Second, I compare the empirical SADs of 226 terrestrial plant communities and 497 mammal communities comprising species of three general trophic guilds (herbivores, omnivores, and carnivores). I analyze the differences in evenness and skewness of the empirical SADs across the different trophic guilds, controlling for species richness, spatial and temporal resolution of the sampling. In model communities, consumer guilds have significantly more even and less skewed SADs than producers. Increasing vulnerability, i.e. the number of predators per prey, also increases consumer evenness and decreases skewness. In empirical datasets, plant communities are significantly less even and more skewed than mammal ones. There are no significant differences in SAD metrics between the different mammal guilds, but carnivores are comparatively rare (i.e. have a higher proportion of species than individuals), whereas omnivores are comparatively more common. Species richness has a positive effect on both evenness and skewness, and spatial and temporal extent have negative effects on evenness and do not affect skewness. I argue that the difference between plant and mammal guilds can be related to higher niche availability in animals than in plants. As no systematic differences were found between the SADs of mammal herbivores, omnivores, and carnivores, this may indicate similar niche availability, when averaged across habitat types, for the different animal trophic guilds.

## Introduction

Most species, in any ecological community, are comparatively rare, and only a few of them are very abundant. This empirical observation is one of the few true universal laws in ecology, a pattern that is observed in all kinds of communities and guilds, from small arthropods (Basset and Kitching, 1991) to tropical trees (He et al., 1997). The Species Abundance Distribution (SAD) describes precisely this variation in species abundances within a given assemblage.

The information contained in SADs, aside from its theoretical interest in understanding community dynamics (Hillebrand et al., 2008), can be of interest for conservation and management (Matthews and Whittaker, 2015) or for estimating ecosystem functions such as primary productivity It is therefore important to understand which are the main axes of variability in SAD shape across guilds, and what are the ecological processes underlying this variability. So far, important efforts have been devoted to understand variation in species abundances across enviromental gradients (Ulrich et al., 2016; Passy, 2016), levels of disturbance (Komonen and Elo, 2017), spatiotemporal scales(Borda-de-Água et al.,, or multiple factors combined (Arellano et al.,2017). Intrinsic differences in species-level traits among the species that make up the community will also be reflected in the SAD. For example, core and satellite species of an ecosystem are likely to differ in their intrinsic abundances regardless of other factors (Magurran and Henderson,2003), and the degree of trophic or habitat specialization of different species may also be related to their relative abundances (Labra et al., 2005; Matthews and Whittaker, 2015).

Despite these many advances, in the debates on the mechanisms that shape SADs, virtually all theories and hypothesis that relate SAD shape to ecological processes at the local scale refer to horizontal communities, i.e. communities of a single trophic level. Thus, these theories emphasize the role of either neutral dispersal and drift (Hubbell, 2001) or horizontal selection (Vellend, 2016). But natural communities form a complex network of species linked by different types of biotic interactions both horizontally (within a given trophic guild) and vertically (across trophic guilds, Fig. **??**). The influence of the interaction topology on the abundance patterns of the different guilds of a community has, to my knowledge, never been explored systematically.

In complex, multitrophic ecological communities, trophic guilds (sensu Fauth et al. 1996) differ in fundamental properties that can potentially be reflected in their associated abundace distributions. First, several studies have demonstrated that basic descriptors, such as biomass or species richness, vary predictably with trophic level. For example, richness pyramids have been shown to be widespread in empirical food webs from different ecosystem and taxonomic types, particularly in marine habitats (Turney and Buddle, 2016). Another distribution, that of biomass across trophic guilds, also has a long history in ecology, starting with the seminal studies by Lindeman (1942) or Odum (1957). Focusing on the relationship between two adjacent trophic levels, Hatton et al. (2015) showed that the biomass of empirical guilds of herbivores and their associated predators scales generally with a power-law of exponent 3/4. They also showed that, in their data, the relationship between mean body mass and community biomass is non-significant for most of the functional groups they studied. As such, if body mass varies in a similar fashion in different trophic levels, the scaling in biomass should also be reflected in a scaling on number of individuals at each trophic level, as foretold by Ramón Margalef (1980).

Other patterns associated to the distribution of abundances, such as species rarity or the degree of dominance, can potentially vary across different trophic guilds, but empirical evidence is scarce. In a study of macroinvertebrate communities, predators showed a higher proportion of species than of individuals in the overall community (Spencer, 2000), pointing to a higher rarity in predator species. Furthermore, Spencer (2000) showed that predator and non-predator species did not vary significantly in the ratios of dominance of the most abundant species. More recently, Dornelas et al. (2011) showed that relative dominance decreased consistently with increasing richness in communities of freshwater fish.

Overall, these lines of evidence suggest a complex, combined influence of the richness and trophic position of a guild on its abundance patterns. However, analyzing abundance distributions in communities comprising several trophic guilds is complicated further by a number of factors. For example, movement capacity generally increases with trophic position (McCann et al., 2005), and in turn, it significantly influences SADs and their variation with sampled area (Borda-de-Água et al., 2017). Therefore, different trophic guilds in the same community will likely require varying sampling areas in order to obtain their abundance patterns in a consistent fashion (see also Holt et.1999). Another issue that needs to be considered when comparing trophic guilds of different communities is precisely how to divide species among guilds in a general way. Broad categories such as herbivores/omnivores/carnivores provide groupings applicable to communities of different ecosystem types, but are likely to be too general, lumping together species with very different ecologies. On the other hand, clearly defined guilds for a given community type (e.g. sap-feeding insects in salt marsh grasses) will be too specific to allow comparisons with guilds from other ecosystem types. Therefore, a robust analysis of the role of trophic guild in SAD metrics needs to control for (1) the variations in richness across guilds, (2) the spatial and temporal extent of the data collection, and (3) the process of defining trophic guilds in a general and informative way.

Here I approach the general question of whether SAD metrics vary predictably across trophic guilds, in two complementary ways. First, I develop a first-principles theoretical approach for studying how different SAD metrics vary in model communities with different degrees of trophic specialization. Second, I analyze SAD patterns of two well-resolved datasets on community abundances of plants (the Gentry forest transects data, Phillips and Miller 2002) and mammals (Thibault et al., 2011). I compare SAD patterns of plant communities and three mammal trophic guilds, controlling for guild richness, spatial and temporal extent of the data collection.

## Methods

### The relationship between SAD properties and trophic level: what does theory predict?

Analyses of species abundance distributions are based, on their first stage, on the grouping of species that are supposed to share certain properties of interest, such as taxonomic relatedness, spatial location or resource use (Fauth et al.1996). As a starting point, I focus on the role of the different trophic guilds in a local multi-trophic community. Intuitively, the variability in number of individuals between species of a given trophic guild will depend on inter-specific variation in both species-level traits and resource use. Assuming, as a working hypothesis, that reproductive strategies, body size and other life-history traits are relatively homogeneous within the species of a given trophic guild, it can be hypothesized that the degree of resource overlap between the species will play an important role in determining their variations in abundance.

This intuitive, qualitative hypothesis can be more precisely formulated by combining network descriptors, body size and numerical abundance patterns, as first proposed by Cohen et al. (2003). These authors derive the resource productivity Π _*j*_ available to species *j* as a simple function of the abundance, body mass and vulnerability of its resource species:

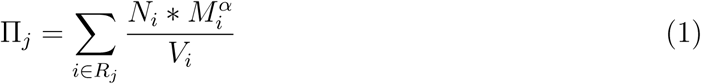

where *R*_*j*_ is the set of resource species for species *j, N*_*i*_ the abundance of resource species *i*, 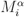 its metabolic rate, and *V*_*j*_ its vulnerability, i.e. its number of predators. Throughout the study, I simplified eq. 1 by setting the scaling coefficient to one to avoid further sources of error, as productivity is thought to vary across guilds (Carbone et al., 2007). The vulnerability of species belonging to a certain trophic guild (assuming that omnivory is negligible) is a measure of the specialization of species from the upper trophic level. Furthermore, the number of individuals of a species, *N*, is a function of its average body mass and the productivity available to it:

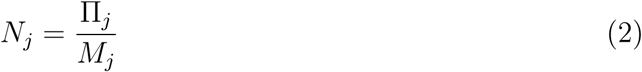

Therefore, with the derivation from Cohen et al. (2003), one can estimate the numerical abundances of species from a given trophic level by knowing their trophic links, average body mass, and the abundances and body masses of their prey. This basic approach may be used as a baseline for predicting the variations of SAD properties with trophic level in closed communities with different patterns of body size variation and/or specialization across trophic levels. The goodness-of-fit of this approximation is analyzed in Appendix 1.

### Metrics for quantifying Species Abundance Distributions

Methodologically, the comparison of SADs is still an unresolved problem in community ecology: there is no standard method for comparing SAD shape of guilds with arbitrary numbers of species or individuals, with most comparisons being qualitative (McGill et al., 2007) or being made between relatively similar communities that don’t differ much in richness or size (e.g. samples from polluted and unpolluted habitats, Matthews and Whittaker 2015). In order to assess the variability between SADs in a general way, robust metrics need to be developed that are independent of number of species and individuals, i.e. that reflect solely the variability in the shape of the distribution. In this study, I assess the variability in SAD shape through two complementary metrics, that quantify the evenness and the skewness of the distribution.

Evenness is defined after the “Hill number” of species diversity, also known as the effective number of species (Jost, 2006). This diversity metric represents how many equally-abundant species would give the observed mean proportional species abundance (Tuomisto, 2012). The evenness metric derived from the effective number of species has a series of desirable properties, summarized in Smith and Wilson (1996). It is also conceptually similar to the variance of a distribution, but the evenness metric has the advantage of having a clear ecological meaning. The skewness metric is simply a robust version of the third moment of a statistical distribution (Brys et al., 2004). Throughout this study, I apply these metrics to the natural abundances of both the simulated and compiled data.

### Variability of SAD metrics with degree of specialization

The framework by Cohen et al. (2003) can be applied to predict how variations in community structure will be reflected in the numerical abundances of consumers, and ultimately, in the shape of the SADs of the different trophic levels. I generated model communities with varying levels of average vulnerability, and estimated the evenness and skewness of the SAD of each trophic level. In particular, I simulated model communities of 100 species divided in four discrete trophic levels, assigning 35 species to the basal trophic level, and 25, 25 and 15 to the upper ones, simulating a slight richness pyramid Turney and Bates.et.al.(2016). Three levels of vulnerability were applied to generate trophic links in which the number of predators is, respectively, 0.1, 0.25, and 0.5 of the species in the upper trophic level. The probability that a link was assigned to any given species is given by its initial abundance, so that more abundant species had a higher degree in the constructed network. Body sizes of the species in the different trophic levels were assigned according to the scaling relationship discussed by Riede et al. (2011). These authors showed that, for a variety of ecosystem types, predator sizes tend to increase with trophic level, but the ratio between predator body mass and prey body mass decreases with trophic level. Equation 1 requires, aside from the vulnerability and body size of each species, the abundances of the basal trophic level to be specified beforehand. I draw basal abundances from a Weibull distribution with scale= 5.6 and shape = 0.9, derived as the average best fit from the 226 sites of the GENTRY dataset (see next section). Variation in these parameters did not alter the qualitative trends observed.

I modelled the response of the SAD metrics of trophic levels 2, 3, and 4 to vulnerability and trophic level with regression models. In particular, I used linear mixed-effect models (R package “lme4”, Bates et al. 2015) with vulnerability and trophic level as fixed effects, and replicate (e.g. the 1000 realizations simulated for each vulnerability level) as a random factor. As I am not interested in predictions from this model, but rather in the effect of the different predictors, I did not perform model selection procedures. However, preliminary analyses showed that more complex statistical models produced qualitatively similar estimates.

### Empirical SAD patterns across trophic guilds

#### Datasets

I analysed two datasets that report species abundances at different sites along with the spatial and temporal effort from each site. Gentry’s Forest Transect Dataset (GENTRY, Phillips and Miller 2002) compiles observations from 226 sites of 0.1 ha in temperate and tropical forests across the globe. At each site, Gentry and collaborators collected the abundance of all plant species with stem diameters equal to or exceeding 2.5 cm diameter at breast height. The second dataset is the Mammal Community Database (MCDB, Thibault et al. 2011), which provides abundances of 660 mammal species in 940 sites, alongside detailed information about the sampling and context of the local community. Rank-abundance curves of one GENTRY site and four MCDB sites are shown in Fig. 2.

**Figure 1:**
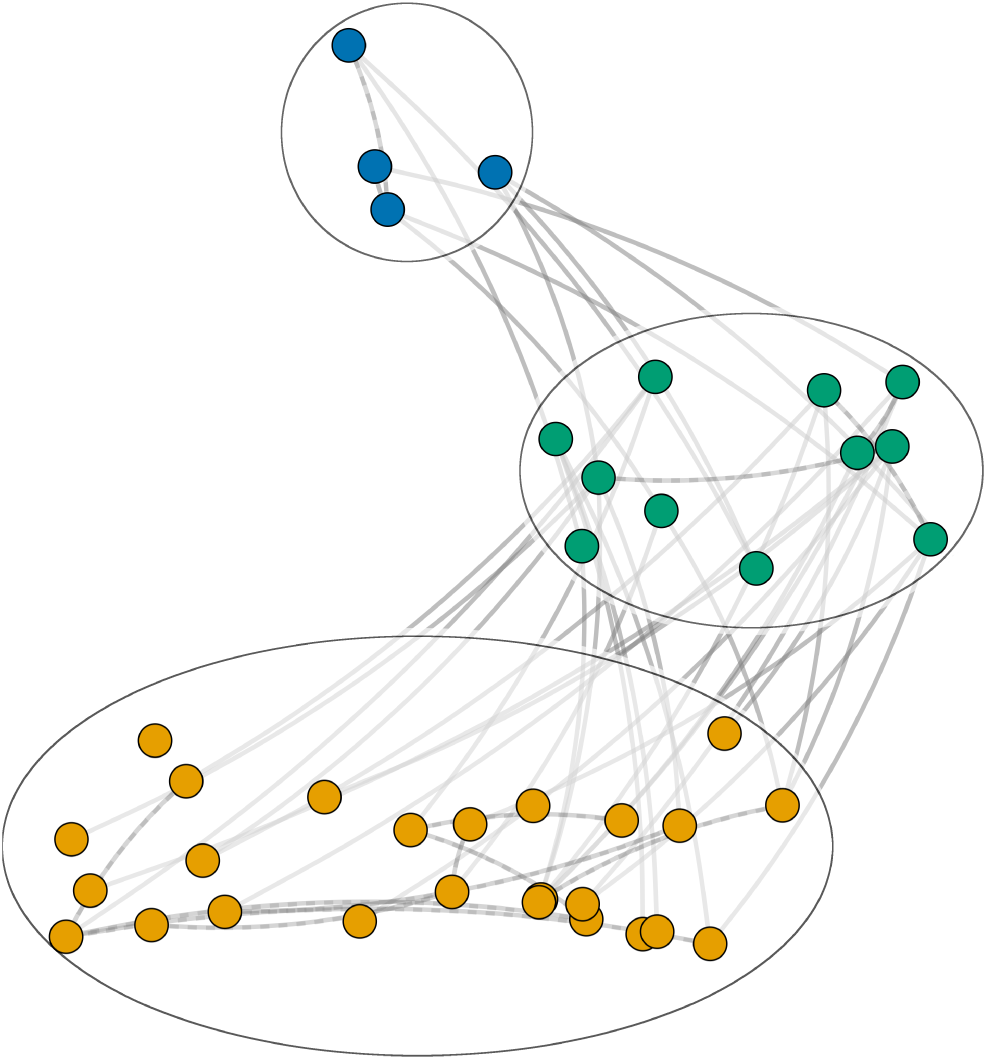
A schematic ecological community with three disjoint trophic guilds. The areas within the ellipses represent the domain of horizontal community ecology that is the subject of most SAD studies, focusing on intra-guild interactions (dashed grey lines) and assuming that interactions with other guilds (full grey lines) are negligible. In this study, I focus on how interactions across guilds drive variations in SAD shape.

**Figure 2:**
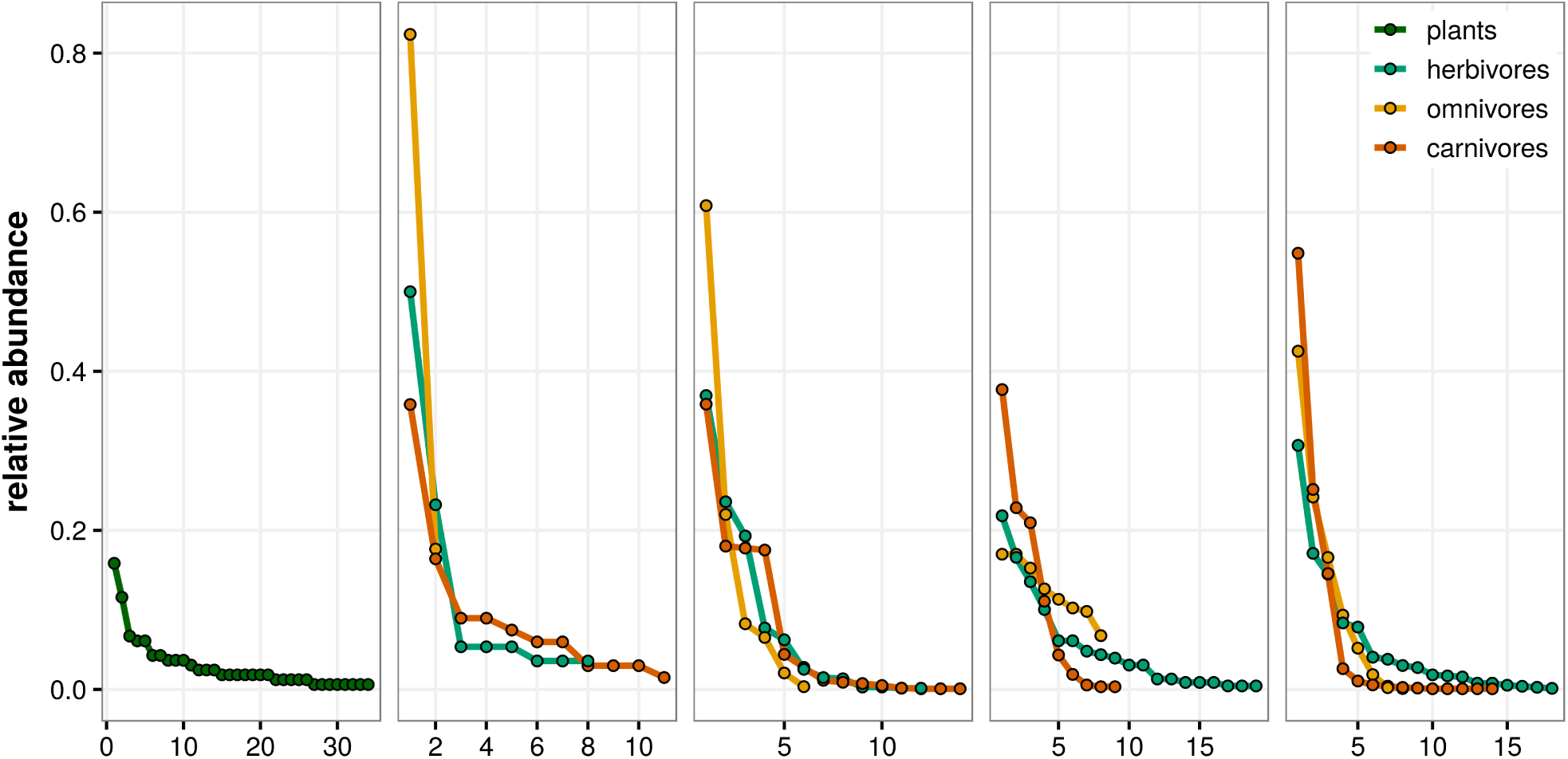
Rank-abundance curves for five sites of the empirical datasets. The leftmost panel shows a curve from a GENTRY site, the other four are sampling sites of the MCDB dataset

Another global dataset on mammal foraging habits (Wilman et al., 2014) was used to derive trophic guild categorizations for every species in the MCDB dataset. I assigned each species to one among three general trophic guilds: (1) herbivores (including folivores, granivores, frugivores, and nectarivores); (2) omnivores, and (3) carnivores (feeding on invertebrates, vertebrates, fish, and carrion). Herbivores and carnivores are species that have at least a 70% of their diet from plant or animal origin, respectively, and species whose diet is < 70% herbivore or carnivore are labelled omnivores. While more detailed trophic guild distinctions are possible, the representativeness of the different categories varies drastically (e.g. there are only 22 mainly frugi/nectarivore species versus 324 foli/granivore ones). In a basic categorization such as the one presented here, the three trophic guilds are well represented (374 herbivores, 145 omnivores, 179 carnivores).

#### Statistical analyses

I calculated the evenness and skewness metrics of each guild at sites in which there is at least three species of that guild, in order to avoid potential biases (for example, a guild of a single species is completely even, or a guild of two species is never skewed). The variation of the SAD metrics with trophic guild was analyzed via statistical models. Evenness values are bounded within the interval [0,1], so a beta regression is an appropriate choice for modelling such bounded data, given the flexibility of the beta distribution. I transformed the evenness values in order to obtain data without proper zeroes and ones, i.e. bounded in (0,1), following Smithson and Verkuilen (2006), and applied a beta regression with trophic guild, species richness, temporal extent and spatial extent as predictors. In particular, I used the R implementation of the GAMLSS family of models (Rigby and Stasinopoulos, 2005), which allows probability density functions to be specified by any number of parameters, themselves functions of the independent variables. I modelled the evenness probability density function with two parameters *μ* and *σ*, named the *location* and *scale* parameters. These parameters are related to the parameterization of the beta distribution in terms of two shape parameters *α* and *β* in the following way:

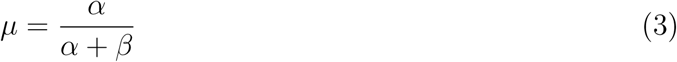

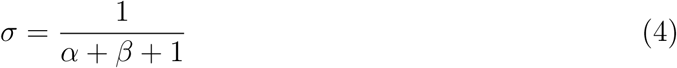

The mean and variance of the beta distribution in terms of *μ* and *σ* is:

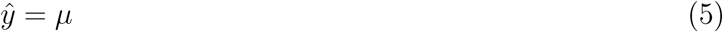

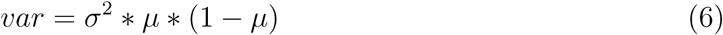

For the final model, I selected the link functions of *μ* and *σ*, and the final set of predictors, via AIC model selection.

Skewness presented a clearly bimodal distribution, with peaks at 0 and 1. Due to the difficulty of modelling such continuous bimodal data, I opted for categorizing the response into three levels of skewness: highly negative, low skewness, and highly positive, represented by the intervals [-1,-0.5), [-0.5,0.5], and (0.5,1]. This response was modelled via a multinomial regression with the same set of predictors as the evenness metric, and AIC model selection was also performed to obtain the final set of predictors.

For the MCDB dataset, it is common to observe species from two or more trophic guilds at the same site (Fig. 2). It is therefore possible to calculate the rarity of each guild as the difference between its proportion of individuals and its proportion of species in the local communities (Spencer, 2000). Furthermore, to complement this calculation, I obtained the relative dominance value of each guild at each site, measured as the abundance of the most abundant species divided by the summed abundances of its guild (Spencer, 2000; Dornelas et al., 2011).

## Results

### Theoretical model

Evenness and skewness metrics show complex, contrasting responses to variations in trophic level and average vulnerability (Fig. 3). After a high increase in evenness from primary producers to consumers, the metric decreases on average with trophic level, whereas for a given trophic level, it increases with increasing vulnerability (Appendix 1: Table 2). The skewness metric again shows an important jump from primary producers to consumers, clearly decreasing. The comparison among the second, third, and fourth trophic level shows a somewhat more complex response: it displays small increases with trophic level, reaching average values close to 0.5 in communities with low or intermediate vulnerability, and moving towards 0 in high vulnerability communities. For a given trophic level, it significantly decreases with increasing vulnerability (Appendix 1: Table 3).

**Table 1:** statistical model coefficients for the evenness of the empirical datasets. See the main text for explanation of the *μ* and *σ* parameters. The *r*^2^ of the overall model is 0.39

| Variable | Estimate | Std. Error | t value | p-value |
| --- | --- | --- | --- | --- |
| **$\mu$ link function: cloglog** |  |  |  |  |
| (Intercept) | -0.509 | 0.039 | -12.824 | < 0.05 |
| trophic.guildherbivores | 0.8681 | 0.055 | 15.882 | < 0.05 |
| trophic.guildomnivores | 0.753 | 0.062 | 11.975 | < 0.05 |
| trophic.guildcarnivores | 0.711 | 0.069 | 10.382 | < 0.05 |
| spatial.extent | $-7.8 * 10^{-7}$ | $3.1 * 10^{-7}$ | -2.481 | < 0.05 |
| temporal.extent | $-3.8 * 10^{-3}$ | $1.3 * 10^{-3}$ | -2.914 | < 0.05 |
| richness | $3.2 * 10^{-3}$ | $2.6 * 10^{-4}$ | 12.126 | < 0.05 |
| **$\sigma$ link function: logit** |  |  |  |  |
| (Intercept) | -0.98 | 0.106 | -9.273 | < 0.05 |
| trophic.guildherbivores | 0.969 | 0.117 | 8.251 | < 0.05 |
| trophic.guildomnivores | 0.461 | 0.135 | 3.407 | < 0.05 |
| trophic.guildcarnivores | 0.819 | 0.143 | 5.709 | < 0.05 |
| spatial.extent | $-1.4 * 10^{-7}$ | $3.9 * 10^{-7}$ | -0.359 | 0.719 |
| temporal.extent | $-6.1 * 10^{-4}$ | $1.5 * 10^{-3}$ | -0.401 | 0.688 |
| richness | $-3.1 * 10^{-3}$ | $8.6 * 10^{-4}$ | -3.624 | < 0.05 |

**Table 2:** statistical model coefficients for the skewness of the empirical datasets. The *r*^2^of the model is 0.15

| Variable | category | Estimate | Std. Error | z score | p-value |
| --- | --- | --- | --- | --- | --- |
| (Intercept) | (0.5,1] | -0.038 | 0.268 | -0.143 | 0.88 |
|  | [-1,-0.5) | -5.773 | 0.688 | -8.39 | < 0.05 |
| trophic.guildherbivores | (0.5,1] | -0.882 | 0.276 | -3.198 | < 0.05 |
|  | [-1,-0.5) | 3.138 | 0.478 | 6.571 | < 0.05 |
| trophic.guildomnivores | (0.5,1] | -1.492 | 0.342 | -4.362 | < 0.05 |
|  | [-1,-0.5) | 2.322 | 0.788 | 2.946 | < 0.05 |
| trophic.guildcarnivores | (0.5,1] | -0.832 | 0.327 | -2.548 | < 0.05 |
|  | [-1,-0.5) | 4.354 | 0.492 | 8.854 | < 0.05 |
| richness | (0.5,1] | 0.012 | 0.003 | 4.427 | < 0.05 |
|  | [-1,-0.5) | -0.314 | 0.224 | -1.397 | 0.16 |

**Figure 3:**
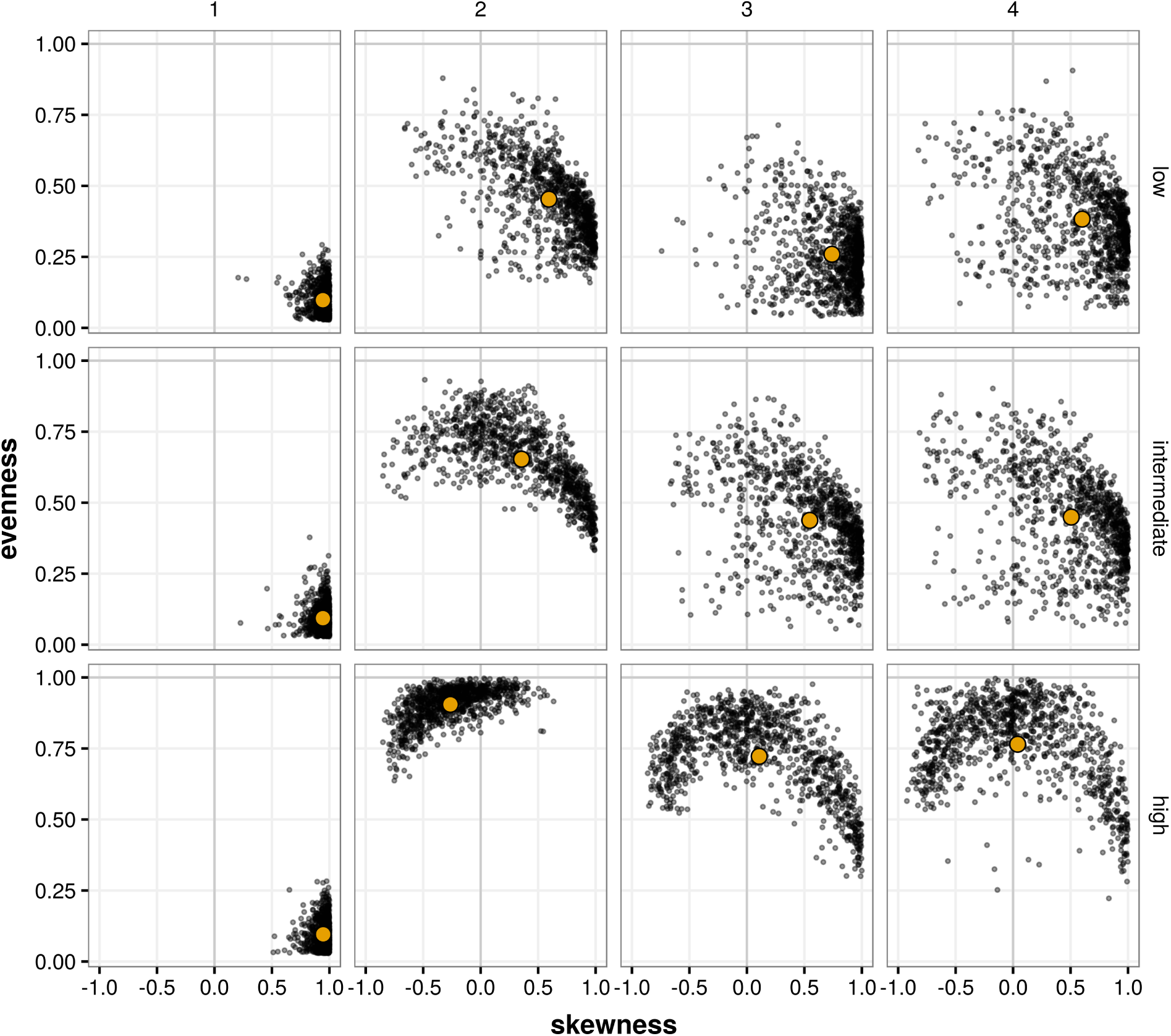
Evenness and skewness values for each combination of trophic level (horizontal panels) and vulnerability (vertical panels) in datasets generated with eq. 1. In the panels, each point represent one of 1000 replicates. The colored point represents the centroid of the two-dimensional distribution.

### Empirical datasets

The statistical model for evenness included all the original predictors (trophic guild, richness, spatial extent and temporal extent, Table 1). Plant guilds were the less even ones overall, showing significant differences with all other guilds (panel (a) of Fig. 4, Appendix 1: Table 4). Among mammal guilds, herbivores show the highest average evenness, after which it further decreases with increasing trophic rank, although the differences are non-significant (Appendix 1: Table 4). This overall pattern of a significant increase from primary producers to consumers followed by a sustained decrease is qualitatively similar to the patterns obtained with eq. 1 (Fig. 3). Richness has a positive effect on evenness, while also decreasing its variability (compare the sign of *μ* and *σ* richness parameters), whereas increasing temporal and spatial extent has a negative effect on mean evenness, and no effect on its variability (Table 1).

**Figure 4:**
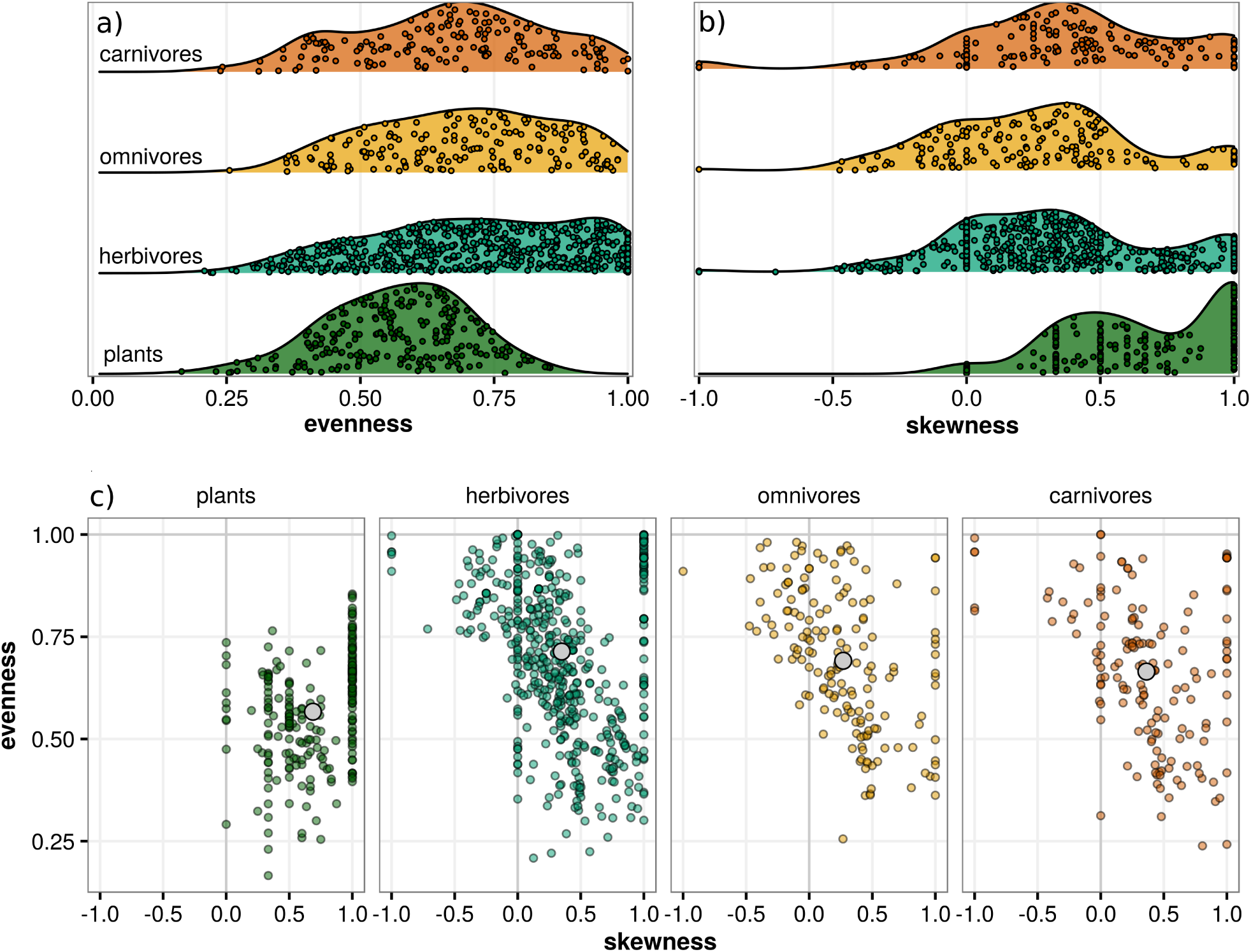
Density distributions of evenness (a) and skewness (b) values of the guilds studied, and the combined distribution of both metrics (panel c, cf. the theoretical results of Fig 3).

The final model for skewness included trophic guild and richness as the only predictors. Again, there were significant differences between plant and mammal guilds in their average skewness (Table 2). Plant and mammal guilds were different mainly when considering low and positive skewness levels, where plants showed the highest average skewness, followed by a drop in all mammal guilds, which showed no statistical differences (Fig. 4, table). The same qualitative pattern is observed in the theoretical model (Fig. 3, particularly the middle panels). The effect of richness is positive, but only significant for the variation between low and highly positive skewness values (Table 2).

The three mammal trophic guilds show similar negative correlations between dominance and richness (panel (a) of Fig. 5, Pearson’s *ρ*: −0.62 for herbivores, −0.66 for omnivores,-0.64 for carnivores). Regarding the relative rarity of the different guilds (panel (b) of Fig. 5), herbivores have the same proportion of species than of individuals in local communities (Wilcoxon signed-rank test, *W* = 131422, *p* = 0.14, Appendix 1: Table 6), whereas omnivores are comparatively more common (i.e. there are higher relative numbers of omnivore individuals than species, *W* = 162226, *p <* 0.05) and carnivores are rarer (there are more carnivore species than individuals, *W* = 18772, *p <* 0.05).

**Figure 5:**
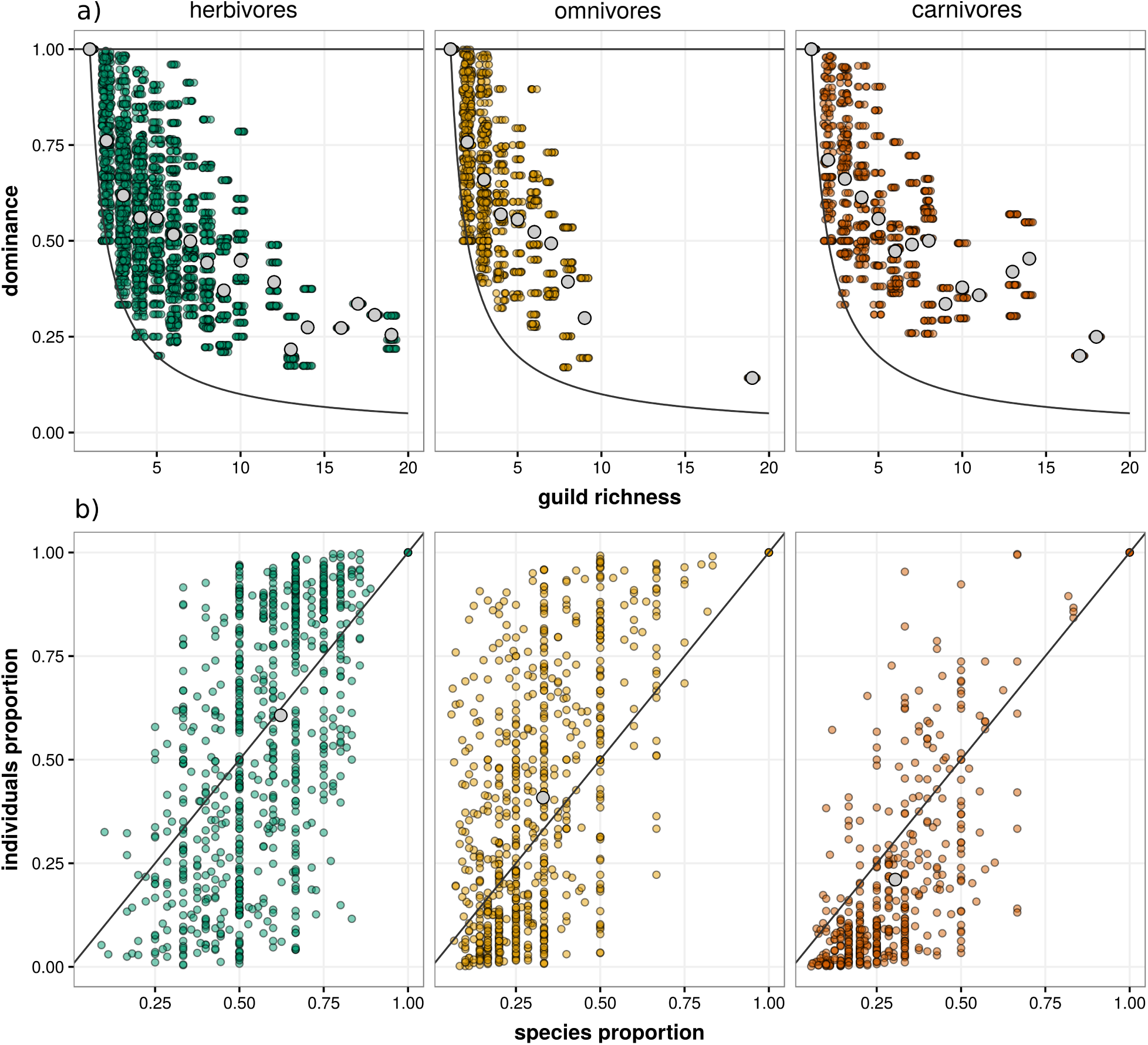
Patterns of dominance, and the proportion of species and individuals of the mammal guilds across sampling sites, where each point is a sampling site. Dominance (upper panel) is defined as the number of individuals of the most abundant species in the guild relative to all individuals from that guild. Minimum and maximum potential dominances are represented by solid curves. The maximum level of richness shown here, for visibility, is 20 species. The lower panel shows the proportion of individuals of a given guild against the proportion of species of the same guild, and the x=y line is shown for visibility. Grey-colored dots in both panels represent average values.

## Discussion

In local communities, the division of species in trophic guilds is informative with regards to the distribution of interactions and biomass flows in the community (Lindeman, 1942; Két al., 2016*b*), but everything else being equal, it is unclear wheter SAD metrics are influenced by the trophic position of a guild in the local community. Here I have shown that there are significant differences between the local SADs of terrestrial plant guilds (primary producers) and different guilds of terrestrial mammals, with plant SADs being less even and more skewed than mammal ones. This result is supported by a first-principles theoretical model. Furthermore, the empirical relationship between SAD metrics and trophic guild is also mediated by species richness and the spatiotemporal extent of the sample.

There are several potential ways of generating theoretical predictions about the variability of SAD metrics with trophic guild. Recent extensions to the theory of island biogeography (Holt, 2009; Gravel et al., 2011) make explicit the differences and feedbacks between species in discrete food chains, and could potentially be extended to predict the variability in species abundances across trophic levels. The model presented here, in turn, is meant to explore the role of local community properties with a fixed number of species and no dynamic migration. In particular, the main question behind it is to explore whether variations in vulnerability (the number of predators of a given prey) influence SAD shape of the increasing trophic levels in communities with a linear trophic structure. I have shown that, given the assumptions of the model, the degree of vulnerability across trophic levels (or species out-degree, in network terminology) is positively related to guild evenness. A high vulnerability in the species of a certain trophic guild implies a high overlap in the trophic interactions of the different predators, something that in this baseline model translates directly to high similartity in net predator abundance, instead of competitive exclusion. One of the main limitations of this model is that vulnerability is assumed constant for all species of the community; it is unlikely that average vulnerability will be maintained across trophic levels, but to the extent of my knowledge, no study has tested this assumption. Another key outcome of the model is that when prey abundance distribution is highly skewed and uneven, the resulting consumer SAD tends to be much more even and less skewed. On the other hand, moderate or high evenness are maintained in higher trophic levels. This tendency to maintain high evenness levels can be due to the above mentioned assumption about equal vulnerability, along with the fact that I did not account for any other species-level trait such as differential assimilation efficiencies, which represent sources of inter-specific variation. Therefore, the model presented here represents very simple closed communities in which basal species abundances determine a static picture of the abundances of species in higher trophic levels. The simplicity of the model is necessary in order to obtain baseline predictions and to uncover the influence of network topology (in the form of vulnerability) on trophic level SADs, without other confounding factors.

Despite the simplicity of the theoretical model, however, its results are qualitatively consistent with the empirical datasets analyzed. The lower evenness and higher skewness observed in terrestrial plant communities relative to mammal ones can be due to a number of factors. Competitive exclusion may be higher in plant communities, leading to higher dominance of a few species, due to the comparatively small set of resources for which plants compete (light, water, and essential nutrients). Mammal guilds, in turn, may potentially have a wider variety of resources available. For example, mammmal herbivores, as classified in this study and according to the *EltonTraits* database (Wilman et al., 2014) used to categorize trophic guilds, can feed mainly on either seeds, fruits, nectar, pollen, and a large list of other plant parts. Therefore, a high degree of specialization within a guild may reduce the importance of competition and, subsequently, the relative differences in abundance between species (Sugihara et al., 2003). Such high specialization in trophic guilds is commonly found in ecological networks (Dunne et al., 2002; Gravel et al., 2011). Other complementary reasons for explaining the variation in SAD metrics between plants and mammals may exist. For example, differences in traits such as movement potential, which can make rare mobile mammal species harder to document, or unavoidable sampling inconsistencies between studies. It may also be the case that plant species are able to maintain lower densities for longer periods than mammal species, thus being more easily observed. Empirical estimates of minimum viable populations indicate that some plant populations may be viable with sizes of < 1000 individuals (Nantel et al., 1996), a number much lower than standard estimates for vertebrates, which range in a few thousands of individuals for their viability (Reed et al., 2003). Therefore, there may simply be more rare plant species than mammals’. Niche differentiation may also be invoked to explain the relative homogeneity in SAD metrics between mammal guilds. As it is the case with herbivores, both omnivores and carnivores may utilize a wide set of resources that can, generally, prevent a high degree of competitive exclusion. All mammal guilds also show a negative correlation between dominance and guild richness, suggesting that richer guilds may be generally more even. The only significant difference between mammal guilds is the relative rarity of carnivore species compared to those of other guilds, which corroborates earlier results by Spencer (2000) on invertebrate communities. These broad-scale results will undoubtedly vary across habitat types, depending on the specific sets of resources available to each guild, and on the local environmental conditions, which may have contrasting effects on the different trophic guilds (Voigt et al., 2003). Aside from potential variations across habitat types or other factors not accounted for in the datasets compiled, I included the effect of species richness and spatiotemporal extent of the sampling in the statistical analyses. In all cases, guild richness was positively associated with evenness, via a decrease in relative dominance, a result in accordance with previous studies (Spencer, 2000; Dornelas et al., 2011). On the other hand, the effects of spatial and temporal extent were negatively related to evenness, which seems counterintuitive given the expected increase in richness with spatial and/or temporal extent of the sampling campaigns. As a first explanation, the richness-evenness relationship is strongest for plant communities, and plant communities in the GENTRY dataset do not vary in either temporal or spatial extent. This could potentially bias the relationships observed here.

All the results presented here assume that trophic interactions are the main driver of variation in species abundances across guilds. In the theoretical results, energy flows from trophic interactions are the only ones considered, and the mammal communities were grouped considering only a general trophic guild classification of the species. This approximation is clearly a simplification of the complex networks of interactions observed in nature. The persistence and abundance of all species in a community is influenced by the whole set of interactions in which they engage, among other factors (Pocock et al., 2012; García -Callejas et al., 2018). However, the distribution and frequency of most non-trophic interactions in empirical communities is not known, so no hypothesis can be formulated at this point regarding their influence on local SADs. In communities in which non-trophic interactions are known, functional guilds can be differentiated by accounting for the set of all interactions in which they engage rather than just trophic ones (Sander et al., 2015; Kéfi et al., 2016*a*); this functional grouping may further reduce intraguild functional variability and thus increase across-guild differences, better reflecting differences in SAD shape across guilds. On the other hand, there is virtually an unlimited number of functional guilds in nature, and establishing a general, cohesive, and manageable set of functional guilds that can be applied to group every potential community seems unfeasible for the time being. Therefore, grouping by trophic guild represents a compromise between generality (as every community can be divided in such manner) and intraguild versus across-guild variability. In any case, the variability in SAD metrics across trophic guilds of a community, and its relationship with other factors, would be better explored with dedicated sampling campaigns of different trophic guilds in multitrophic communities, in which properties such as vulnerability can also be tracked. This could be carried out, for example, in pond mesocosms in which most species are confined to the pond habitat, or for terrestrial habitats, in small islands where complete censuses of different trophic guilds are feasible.

The existence of an intrinsic variability in SAD shape across functional guilds has important consequences for both fundamental and applied community ecology. In theoretical models, the contributions of intraguild and interguild interactions need to be integrated in a general mechanistic framework, in order for their relative importance to be estimated. This shift from intraguild, competitive interactions to multi-trophic, community scale thinking is a necessary step forward in theoretical community ecology (Chesson and Kuang, 2008; Godoy et al., 2018; Seibold et al., 2018). Empirical analyses of SADs would also benefit from incorporating information about the communities that harbor the guilds under study. In studies analyzing intraguild drivers of SAD shape, the community context of the study should be used to establish null expectations of interguild influence, for example, based on the trophic relationships with other guilds, as in eq. 1. If the guilds under study are not functionally homogeneous (e.g. taxonomic assemblages sensu Fauth et al. 1996) deriving mechanistic explanations about SAD shape is usually not possible, due to interspecific differences in resource use, trophic position, etc. In such descriptive studies, therefore, the community context is unnecessary. Meta-analyses of Species Abundance Distributions have shown great disparity regarding the most appropriate statistical models for fitting SADs (Ulrich and Gotelli, 2010; Baldridge et al., 2016). If the results presented here hold, accounting for this null expectation may help clarify which statistical models are best suited to fit each set of data.

## Conclusions

Species Abundance Distributions have many axes of variability. Here I have showed that intrinsic differences exist between the SAD of terrestrial plant and mammal communities. Plant communities are significantly less even and more skewed than mammal ones, and there are no significant differences in either metric between the mammal trophic guilds considered. This result may arise from differences in niche availability for the different guilds, following the hypothesis that higher niche availability implies a higher evenness in species abundances. Although these results are derived from extensive datasets controlling for several factors, targeted studies are needed to further confirm this pattern and test it in a variety of systems. This prospective line of research would shed new light on both theoretical and applied analyses of Species Abundance Distributions, and would help in the integration of classic horizontal community ecology patterns in the context of communities encompassing multiple trophic levels and interaction types.

## Data availability

GENTRY, MCDB, and EltonTraits datasets are publicly available (see references). All the code used to generate the results is available at the repository of the study: https://github.com/DavidGarciaCallejas/SAD

