## Supplementary Materials for "On the variability of Species Abundance Distributions with trophic guild and community structure" (from another version of this preprint: version 1, 2018-03-27)

Appendix A: Supplementary tables and figures

**Table A1**: estimated regression parameters for the evenness metric in the simulated datasets.

| | estimate | std. error | t value | p-value |
|---|---|---|---|---|
| **$\mu$ link function: logit** | | | | |
| Intercept | 1.122 | 0.005 | 202.507 | $< 0.05$ |
| VulnerabilityIntermediate | 2.379 | 0.009 | 247.407 | $< 0.05$ |
| VulnerabilityHigh | 3.374 | 0.013 | 252.686 | $< 0.05$ |
| trophic.level3 | 0.069 | 0.007 | 9.108 | $< 0.05$ |
| trophic.level4 | 0.637 | 0.008 | 76.566 | $< 0.05$ |
| **$\sigma$ link function: log** | | | | |
| Intercept | 5.027 | 0.014 | 356.3 | $< 0.05$ |
| **$\nu$ link function: logit** | | | | |
| Intercept | -25.39 | 1054.09 | -0.024 | 0.98 |

**table A2:** estimated regression parameters for the skewness metric in the simulated datasets. The estimated $\sigma_{replicate}$ is 0.032

| | estimate | std. error | t value | p-value |
|---|---|---|---|---|
| Intercept | $-9.09 * 10^{-2}$ | $5.66 * 10^{-3}$ | -16.067 | $< 0.05$ |
| VulnerabilityIntermediate | $-1.62 * 10^{-2}$ | $6.11 * 10^{-3}$ | -2.65 | $< 0.05$ |
| VulnerabilityHigh | $1.97 * 10^{-2}$ | $6.11 * 10^{-3}$ | 3.157 | $< 0.05$ |
| trophic.level3 | $8.59 * 10^{-3}$ | $6.11 * 10^{-3}$ | 1.407 | 0.16 |
| trophic.level4 | $3.62 * 10^{-3}$ | $6.11 * 10^{-3}$ | 0.593 | 0.55 |

**table A3:** estimated regression parameters for the evenness metric in the empirical datasets. The estimated $\sigma_{taxon}$ is 0.475

| | estimate | std. error | t value | p-value |
|---|---|---|---|---|
| **$\mu$ link function: logit** | | | | |
| Intercept | 0.812 | 0.048 | 17.039 | $< 0.05$ |
| trophic.guildDetritivore | 0.065 | 0.061 | 1.077 | 0.28 |
| trophic.guildOmnivore | 0.059 | 0.053 | 1.109 | 0.27 |
| trophic.guildCarnivore | 0.359 | 0.047 | 7.62 | $< 0.05$ |
| habitatFreshwater | -0.08 | 0.043 | -1.854 | 0.06 |
| habitatTerrestrial | -0.101 | 0.044 | -2.287 | $< 0.05$ |
| **$\sigma$ link function: log** | | | | |
| Intercept | 1.197 | 0.023 | 51.38 | $< 0.05$ |
| **$\nu$ link function: logit** | | | | |
| Intercept | -1.077 | 0.035 | -30.72 | $< 0.05$ |

**table A4:** estimated regression parameters for the skewness metric in the empirical datasets. The estimated $\sigma_{taxon}$ is 0.714

| | estimate | std. error | z value | p-value |
|---|---|---|---|---|
| **Location coefficients** | | | | |
| trophic.guildDetritivore | -0.131 | 0.186 | -0.702 | 0.48 |
| trophic.guildOmnivore | -0.372 | 0.155 | -2.390 | $< 0.05$ |
| trophic.guildCarnivore | -0.577 | 0.162 | -3.562 | $< 0.05$ |
| habitatFreshwater | 0.033 | 0.130 | 0.253 | 0.80 |
| habitatTerrestrial | 0.021 | 0.326 | 0.065 | 0.94 |
| **Threshold coefficients** | | | | |
| $< 0.5$ \| [-0.5,0.5] | -4.757 | 0.412 | -11.545 | - |
| [-0.5,0.5] \| 0.5 | 1.798 | 0.390 | 4.610 | - |

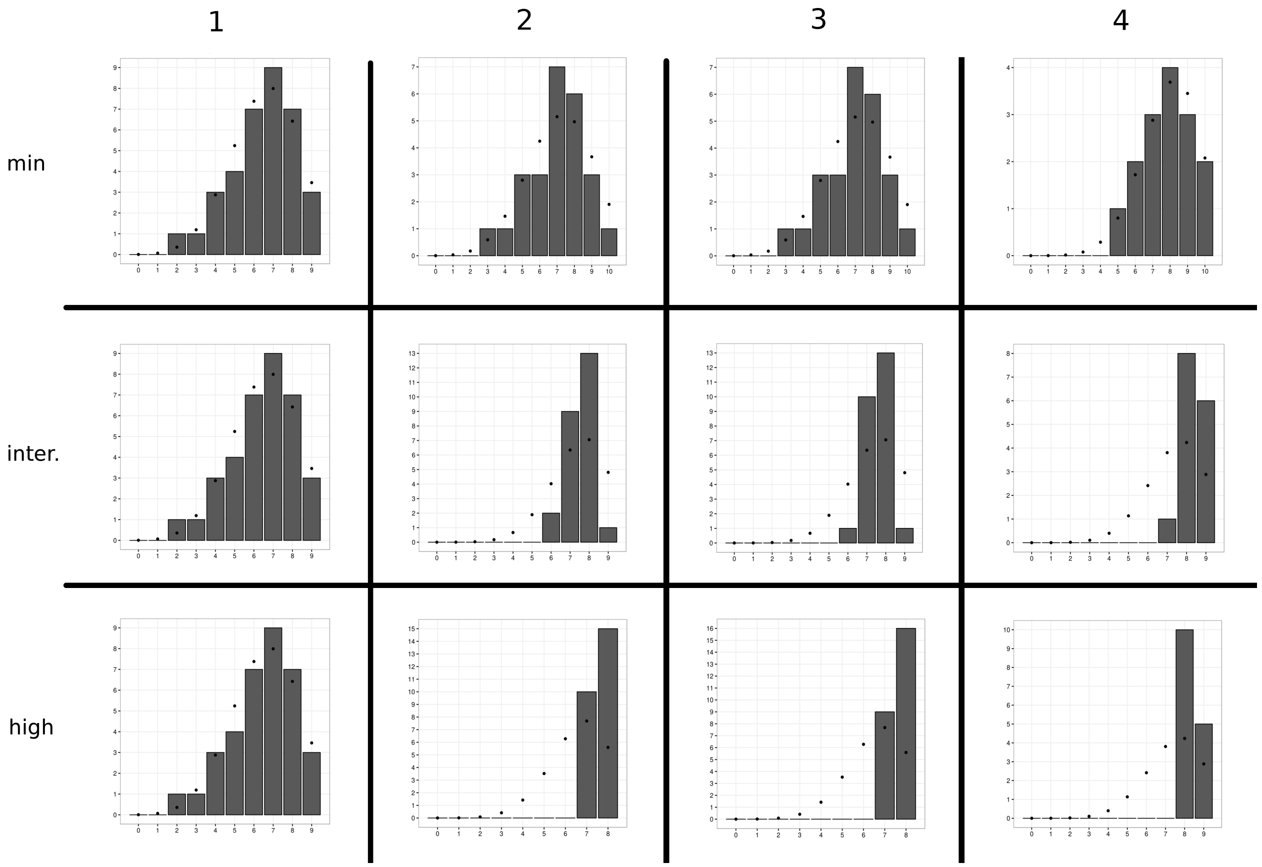


Figure 1: Species Abundance Distributions of model datasets, arranged by trophic level (1-4) and vulnerability level (minimum, intermediate, high). The SADs depicted represent the average SAD over 1000 replicates. SADs are fitted to a gambin distribution (black dots) for reference.

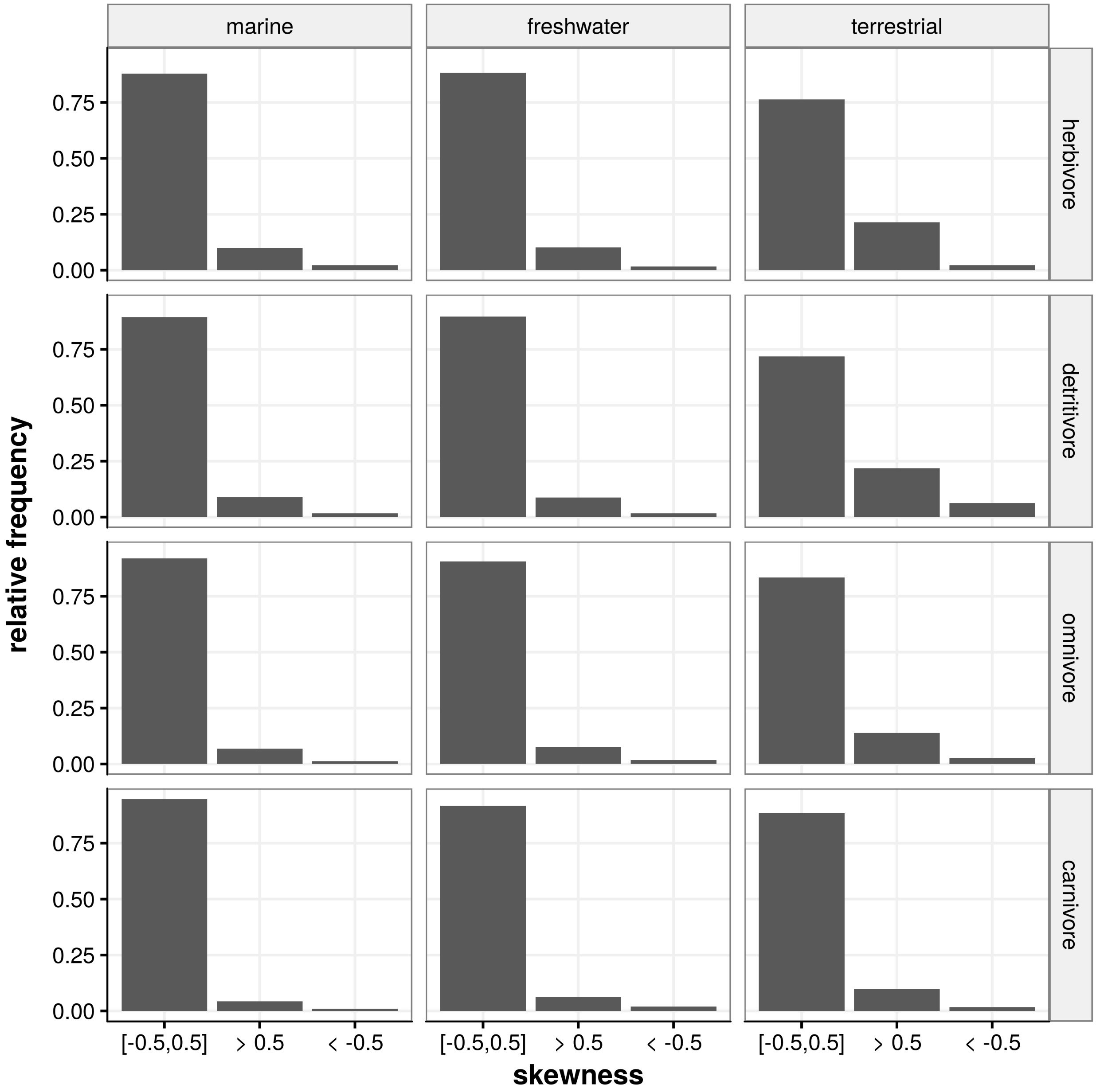


Figure 2: Skewness levels observed in empirical datasets. Bars represent the relative frequency of each skewness level for each combination of trophic guild and habitat type.
